# Hypervirulent pneumococci display high levels of nasopharyngeal shedding and rapid onward transmission

**DOI:** 10.1101/2023.06.17.545125

**Authors:** Murielle Baltazar, Laura C. Jacques, Teerawit Audshasai, Marie Yang, Aras Kadioglu

## Abstract

*Streptococcus pneumoniae* serotype 1 is a major cause of invasive pneumococcal disease. Despite its high attack rate, serotype 1 exhibits a low carriage prevalence within the population, which raises important questions about the relationship between carriage and transmission of hypervirulent pneumococcal strains between individuals. We compared the transmission dynamics of serotype 1 sequence type ST217 to serotype 2 strain D39 using a novel model of transmission in young adult mice. Donor index mice were intranasally infected with ST217, D39 or isogenic pneumolysin-deficient mutants and co-housed with recipient naive contact mice. Three days later, all mice were infected with influenza A virus (IAV). Pneumococcal transmission was analysed during colonisation alone and co-infection with IAV by quantification of shedding and nasal colonisation in index and contact mice. The role of the toxin pneumolysin in shedding, transmission and colonisation, and the host nasopharyngeal immune response were investigated. We show that ST217 was shed in index mice at significantly greater levels compared to D39. Upon viral co-infection, ST217 was shed and transmitted at a faster rate to contact mice and displayed higher transmission levels compared to D39. Interestingly, the toxin pneumolysin did not play a role in shedding. However, upon acquisition by contact mice, pneumolysin-dependent macrophage recruitment was observed in the nasopharynx. Our results show that the rapid and high transmission rate of serotype 1 is a key factor in its ability to disseminate quickly within the population and cause disease outbreaks.

## Introduction

*Streptococcus pneumoniae* (the pneumococcus) is a major human pathogen responsible for invasive pneumococcal diseases (IPD) such as pneumonia, meningitis and bacteraemia. Colonising the human nasopharynx, the pneumococcus can establish a commensal relationship with its host leading to a state of “carriage” which is asymptomatic in the majority of cases (1–3). However, the pneumococcus can also spread into neighbouring tissue compartments and cause life-threatening invasive diseases, particularly in the very young, the elderly and immunodeficient patients (4, 5).

Serotype 1 is one of the most prevalent invasive pneumococcal serotypes in West and Sub-Saharan Africa, which includes the sequence type (ST) 217 hypervirulent clonal complex (6–11). Interestingly, despite a very low prevalence in healthy carriers (< 1-2%) (12) and a short carriage duration in humans, estimated at approximatively 9 days before clearance (13), serotype 1 is a major cause of IPD outbreaks (8, 14–16), mainly due to transmission in older children, adolescents and adults (17), suggesting that its acquisition and transmission is more transient and dynamic than other serotypes. However, the mechanisms behind serotype 1 transmission between individuals remain unclear.

Pneumococcal carriage which is the first stage for human-to-human transmission, usually occurs through shedding of pneumococci in respiratory droplets spread from colonised individuals (2, 18, 19). Fomites contaminated with oropharyngeal secretions containing pneumococci can also act as the starting point for transmission (20, 21). Studies have shown that pneumococcal transmission occurs more frequently during cold and dry months (22–24) and during periods when nasal discharge is prevalent; a common symptom during co-infection of the upper respiratory tract with respiratory viruses (25–28). On the other hand, epidemiological studies have shown that adults can act as reservoirs for transmission, leading to subsequent outbreaks of infection within the larger population, particularly with highly virulent serotypes such as serotypes 1 or 3 (9, 29, 30).

To date, only a ferret model (31) and two neonate/infant transmission models in mice and rats respectively (32, 33) have been developed to study pneumococcal transmission. Both rodent models use neonate/infant animals which have immature immune systems and are useful for modelling equivalent responses in human infants, but are less suitable for modelling older children, adolescents and adults who have developed immune systems. To date, no adult rodent model has been developed to investigate the mechanisms driving pneumococcal transmission in immunologically mature/competent hosts. We sought to address this absence of knowledge by developing a mouse to mouse transmission model using young mice equivalent to human children/adolescents in age range.

The aim of this study was to investigate the relationship between carriage and transmission dynamics of a hypervirulent serotype 1 ST217 clinical isolate in comparison to the well characterised serotype 2 (strain D39) pneumococcus, and to determine how transmission affects the newly colonised host by developing a clinically relevant novel mouse model of transmission using young adult mice. In addition, we investigated the role of the major pneumococcal virulence factor, the pneumolysin, in transmission and the immune response of the new host upon pneumococcal acquisition.

## Methods

### Ethics statement

This study was conducted in strict accordance with the guidelines outlined by the UK Home Office under the Project License Number PB6DE83DA. All animal experimentations were performed at the Biomedical Services Unit, University of Liverpool and approved by the University of Liverpool Animal Welfare and Ethical Review Body.

### Pneumococcal and viral strains and culture conditions

*S. pneumoniae* serotype 1 ST217 and serotype 2 strain D39 (NCTC 7466), and their isogenic pneumolysin-deficient mutants are listed in supplementary table S1. Challenge doses were prepared as previously described (34). Influenza virus A/HKx31 (X31, H3N2) was propagated in the allantoic cavity of 9-day-old embryonated chicken eggs at 35°C for 72 h. Viral titres of infection doses and in nasopharyngeal mouse tissues were determined by plaque assay using MDCK cells with an Avicel overlay.

### Construction of pneumolysin-deficient serotype 1 ST217 mutant

The isogenic serotype 1 ST217 pneumolysin-deficient (ST217*Δply*) mutant was constructed by replacing the *ply* gene with the *aphA3* gene (conferring resistance to kanamycin) through allelic replacement. A ST217*Δply* mutant was selected on selective medium and verified by PCR and Sanger sequencing.

### Pneumococcal transmission in young mouse model

Young adult female CD1 mice were obtained from Charles River Laboratories (Kent, UK). Upon delivery, mice were allowed to acclimatise for seven days prior to use. The transmission model described in this study and schematically represented in Figure 1A, is a modification of a previous described study (32). A 2:3 ratio of colonised index mice to naïve contact mice was used. On day zero, index mice were intranasally infected with *S. pneumoniae* and co-housed in the same cage with naïve contact mice for three days. On day 3 post infection, all mice were intranasally infected with influenza virus A/HKx31 (X31, H3N2). Pneumococcal shedding and transmission from index mice to contact mice was quantify by culture of the nasal secretions collected from nose tapping from day 1 to day 9 post infection. Mice were euthanised on day 10 post infection and nasopharyngeal tissues were collected to quantify the level of colonisation.

**Figure 1.**
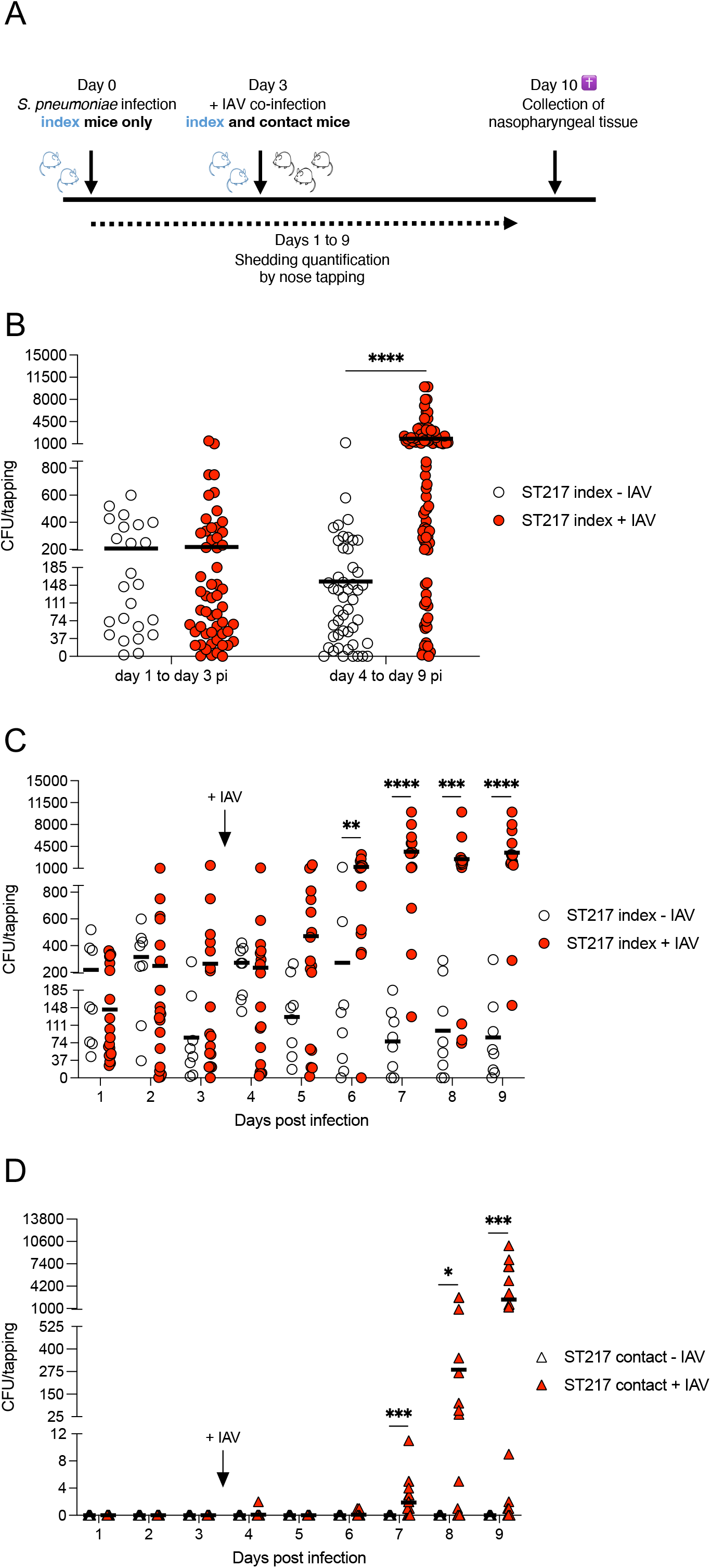
Serotype 1 ST217-shedding at high level leads to its transmission. (*A*) Timeline of the course of the experiment. On day zero, *S. pneumoniae* was intranasally administered to young adult index mice. Three days post pneumococcal infection, all index and contact mice were intranasally infected with influenza virus A/HKx31 (X31, H3N2) (+ IAV) or mock inoculated with PBS. Pneumococcal daily shedding was collected and quantified from culture of nasal secretions from day 1 to day 9 post infection. On day 10, all mice were euthanised, nasopharyngeal tissues were collected, and pneumococcal colonisation density was quantified. (*B*) Comparison of levels of shedding of serotype 1 ST217 in absence (from day 1 to day 3 post infection) and presence (from day 4 to day 9 post infection) of IAV. Comparison of daily shedding in index (*C*) and contact (*D*) mice over time. Each symbol represents CFU/tapping counted from a single mouse on one day and the mean value is indicated. Shedding on day 3 was quantified prior to IAV infection. Statistical differences between the two groups in each day were determined using the Mann-Whitney *U* test. *P < 0.05, ** P < 0.01, *** P < 0.001 and **** P < 0.0001. Data represent results of multiple independent experiments (n = 1-5).

### Measurement of the capsule thickness

The pneumococcal capsule thickness was determined by measuring the zone of exclusion of FITC-dextran based on a modification of the method of Gates et al (35) using FITC-dextran (2000 kDa, Sigma).

### Immune response analysis by flow cytometry

Flow cytometry was performed on cells from nasopharyngeal tissues collected from mice using a BD FACSCanto™ II flow cytometer (BD Biosciences). Data were analysed with FlowJo™ software, version 10.6.2 (BD). All monoclonal antibodies used for cell staining are listed in supplementary table S2.

### Statistical analysis

Statistical analysis was performed using GraphPad Prism 8^®^ (GraphPad Software, Inc., USA). Differences were determined using the Mann-Whitney *U* test, the Kruskal-Wallis test with Dunn’s post test or the Fisher’s exact test for comparing respectively, two groups, multiple groups and proportions.

## Results

### High level of shedding is required for transmission in young mice

We first investigated transmission of serotype 1 ST217 in young mice in the absence and presence of IAV co-infection, whereby viral infection has been shown to facilitate pneumococcal transmission (32, 36). On day zero, (donor) **index** mice were intranasally infected with ST217 and co-housed with naïve (recipient) **contact** mice. The challenge dose given allowed for stable nasopharyngeal colonisation without any seeding into lungs or blood (37). Three days after pneumococcal infection, either all mice (both index and contact) were intranasally co-infected with IAV or they were mock inoculated with PBS (Figure 1A).

Pneumococcal shedding quantified by nose tapping over 9 days showed that between days 1 and 3 post pneumococcal infection (prior to IAV co-infection), index mice from both groups shed bacteria at equivalent levels (Figure 1B). Following infection with IAV from day 4 to day 9 post infection, numbers of bacteria shed by index mice increased significantly compared to mono-infected index mice for which ∼ 12-fold less bacteria were shed (p < 0.0001) (Figure 1B). Specifically, the level of ST217-shedding was significantly greater 72 h following IAV infection and remained significantly higher over time compared to index mice colonised with ST217 only (Figure 1C). Consequently, ST217 transmission from index to contact mice occurred mostly from day 7 onwards (96 h post IAV infection) (Figure 1D), suggesting that high shedding was required in donor individuals to induce transmission in co-housed recipient mice.

Enumeration of CFU in total nasopharynx at day 10 post infection showed that all index mice were colonised with ST217, although IAV co-infection led to significantly increased density of pneumococci (∼ 100-fold more) compared to infection with pneumococcus alone (p < 0.0001) (Supplementary figure S1). These results indicate that the presence of influenza virus correlated with significant increases of nasal pneumococcal density in young donor mice which significantly increased shedding of serotype 1 ST217 leading to its transmission.

### Serotype 1 displays high levels of shedding leading to high rates of transmission

We next compared kinetics of serotype 1 ST217 shedding and transmission to serotype 2 strain D39. Prior to IAV co-infection, shedding events were detected in D39-colonised index mice, albeit at a significantly lower level compared to ST217 (p < 0.0001) (Figure 2A). Upon IAV co-infection however, D39-shedding was greatly increased in index mice, although again, at a significantly lower level compared to ST217 (Figures 2B and 2C). Viral infection also enabled successful transmission of D39, as shedding events were detected in contact mice, albeit at a significantly lower level (p < 0.01) (Figure 2B). ST217-shedding events were detected as early as days 4 and 6 while D39 was shed from day 7 (Figure 2D), suggesting that successful transmission of ST217 occurred at a faster rate than D39.

**Figure 2.**
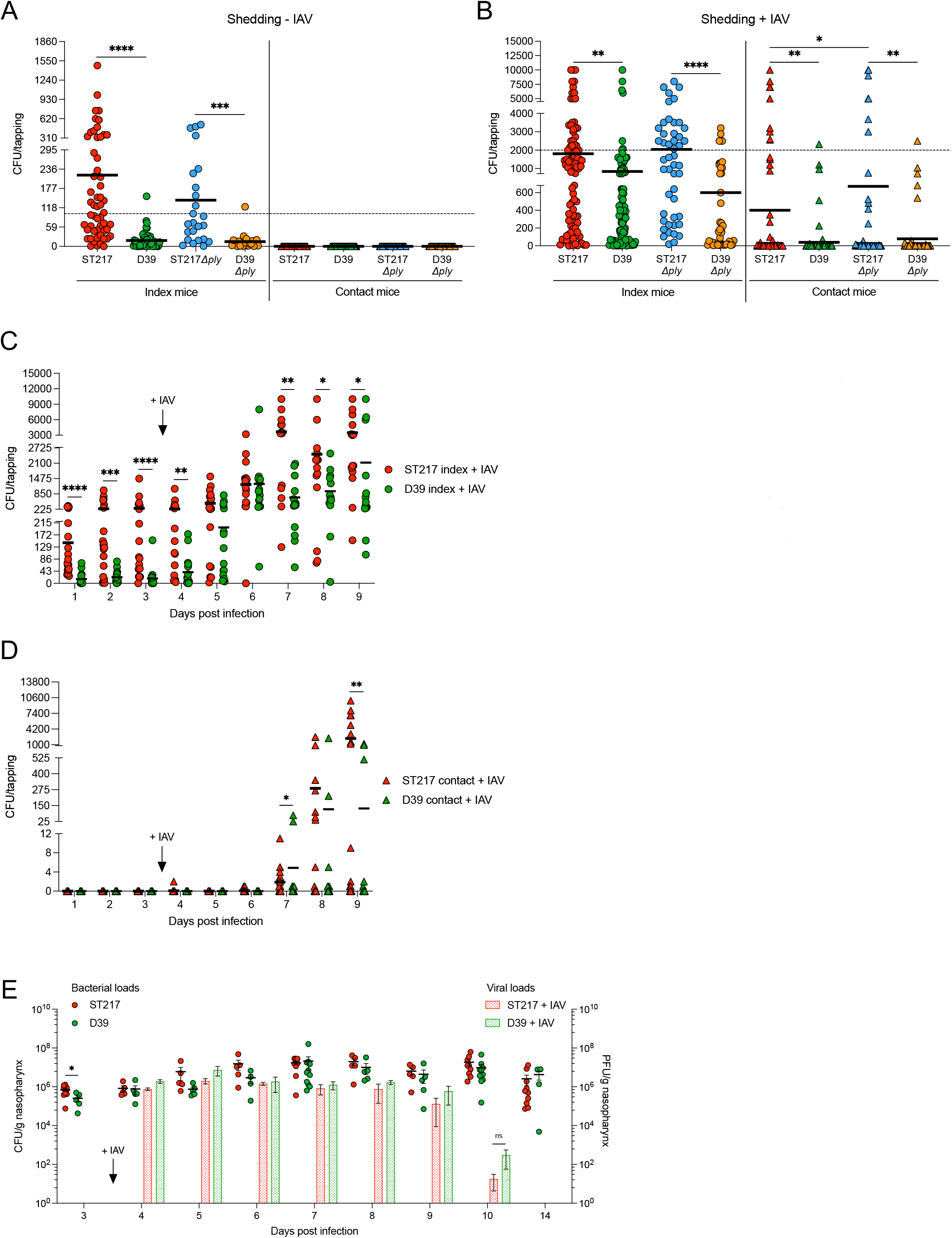
Serotype 1 ST217 displays high levels of shedding. On day zero, *S. pneumoniae* was intranasally administered to young adult index mice. Three days post pneumococcal infection, all index and contact mice were intranasally infected with influenza virus A/HKx31 (X31, H3N2) (+ IAV). Pneumococcal daily shedding was collected and quantified from culture of nasal secretions from day 1 to day 9 post infection. Comparison of levels of shedding of serotype 1 ST217, serotype 2 (D39) and the isogenic *Δply* mutants in absence (from day 1 to day 3 post infection) (*A*) and presence (from day 4 to day 9 post infection) (*B*) of IAV. Comparison of daily shedding in index (*C*) and contact (*D*) mice over time. (*E*) Comparison of bacterial and viral loads in nasopharynx of mice infected with serotype 1 ST217 or serotype 2 (D39) in absence (day 3 post infection) and presence (from day 4 to day 14 post infection) of IAV. Shedding on day 3 was determined prior to IAV infection. The black dotted panel-wide horizontal lines represent respectively the 100 CFU and 2,000 CFU threshold levels described in Results. Each symbol represents CFU/tapping or CFU/g and PFU/g nasopharynx counted from a single mouse on one day and the mean (*A-D*) or mean ± SEM (*E*) values are indicated. Statistical analysis was performed using the Kruskal-Wallis test with Dunn’s post test (comparing multiple groups) or the Mann-Whitney *U* test (comparing two groups in each day). *P < 0.05, **P < 0.01, ***P < 0.001 and ****P < 0.0001. ns: not significant. Data represent results of multiple independent experiments (n = 1-5).

Despite clear differences in ST217 and D39 shedding and transmission following IAV co-infection, there were no significant differences in overall nasopharyngeal bacterial and viral loads between the two groups of mice over a 14-day colonisation period (Figure 2E), suggesting that bacterial or viral infection loads *per se* were not the instigator of increased ST217 shedding and transmission over D39.

Interestingly, despite near equivalent nasopharyngeal CFU density, greater proportions of “high-shedding” events were observed in index mice shedding ST217. Indeed, prior to IAV co-infection, 54% of ST217-shedding events were greater than 100 CFU/tapping compared to D39 for which only 2% of shedding events exceeded this threshold (p < 0.0001), while post IAV co-infection, 30% of ST217-shedding events were greater than 2,000 CFU/tapping compared to just 8% for D39 (p < 0.001) (Supplementary figure S2A). The proportion of contact mice shedding bacteria was also significantly higher following ST217-acquisition (92%) compared to D39-acquisition (43%) (p < 0.001) (Supplementary figure S2B), clearly demonstrating that ST217 had a significantly greater propensity to be shed and at higher levels than D39.

No significant difference in nasopharyngeal bacterial density was observed between ST217- and D39-infected index mice at day 10 post infection (Figure 3A). However, contact mice tended to have a higher density of ST217-pneumococci compared to contact mice that acquired D39 (p = 0.0501) (Figure 3A). A significantly higher transmission rate of 58% was observed with ST217 compared to D39 which colonised only 24% of contact mice (p < 0.05) (Figure 3B), suggesting that serotype 1 has a higher transmission rate than serotype 2 (D39).

**Figure 3.**
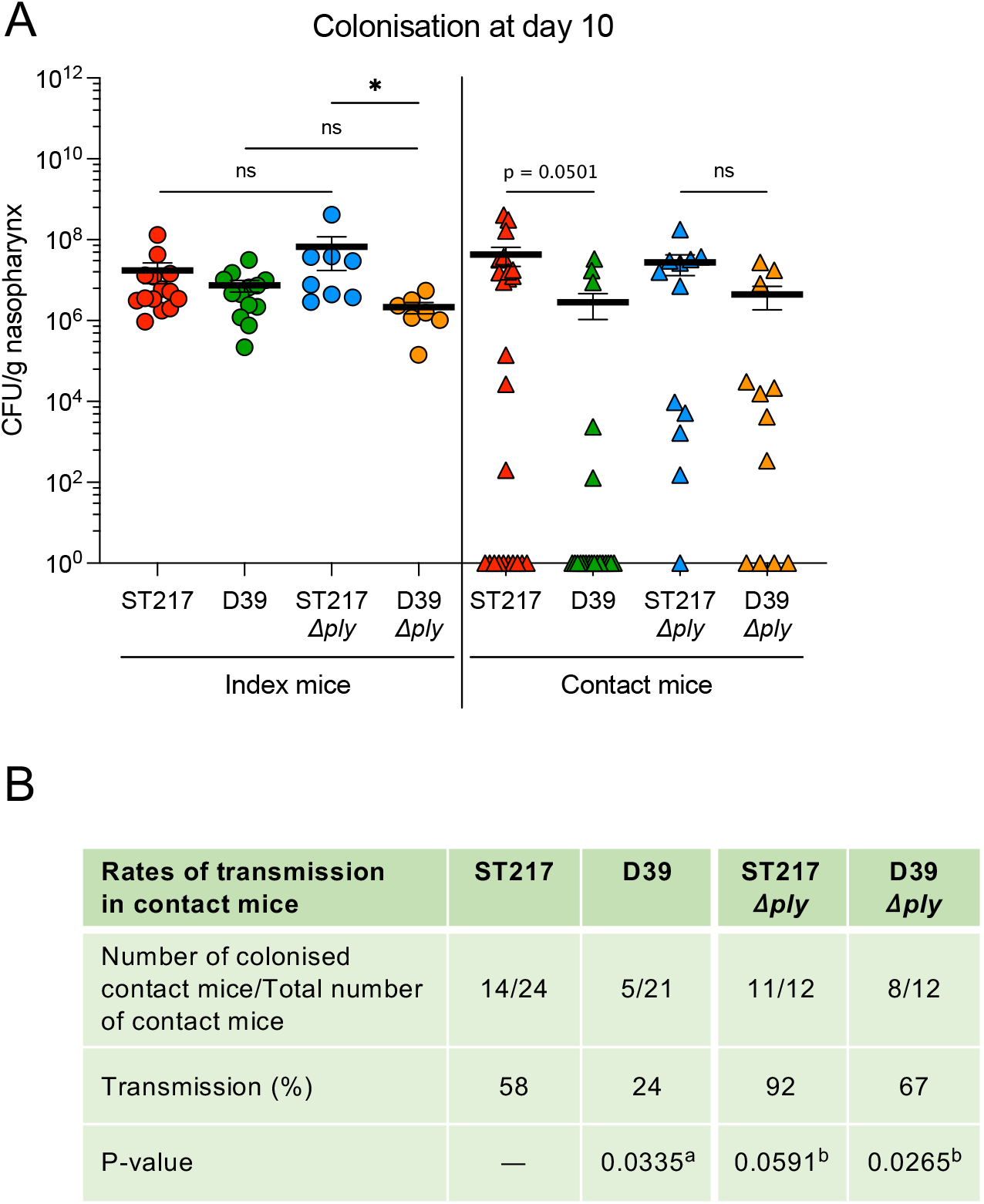
Serotype 1 ST217 displays a high rate of transmission independently of pneumolysin. (*A*) Colonisation densities of serotype 1 ST217, serotype 2 (D39) and the isogenic *Δply* mutants in the nasopharynx of both index and contact mice at day 10 post infection during co-infection with IAV. Each symbol represents CFU counted from a single mouse and the mean ± SEM value is indicated. (*B*) Rates of pneumococcal transmission in contact mice. Statistical analysis was performed using the Kruskal-Wallis test with Dunn’s post test (comparing multiple groups) or the Fisher’s exact test (comparing proportions). *P < 0.05, ns: non significant. ^a^ in comparison to ST217; ^b^ in comparison to wild-type strains. Data represent results of multiple independent experiments (n = 1-5).

The expression of large amounts of polysaccharide/thick capsule has been shown to promote shedding and transmission (38). In vitro comparison of the capsule size of both strains showed that ST217 had a significantly thicker capsule than D39 (p < 0.0001) (Supplementary figure S3), suggesting that a thicker capsule may have increased its transmissibility to recipient individuals.

### Pneumolysin deficiency promotes nasal colonisation upon acquisition in the new host and pneumococcal dissemination within population

We sought to determine the role of the major virulence factor pneumolysin in shedding and transmission. We constructed an isogenic knockout mutant in serotype 1 ST217 in which the *ply* gene was completely deleted and replaced by a linear DNA containing the *aphA3* gene conferring resistance to kanamycin. Shedding and transmission dynamics of the pneumolysin-deficient mutants were then compared to those of their respective parent strains.

In the absence of IAV, no significant differences were observed in index mice (Figure 2A). Upon IAV co-infection, increases in levels of shedding were observed, however, both *Δply* mutants were still shed at equivalent levels to their parent strains (Figure 2B). As observed with wild-type isolates, highly significant differences in shedding were observed in index mice infected with the ST217*Δply* mutant compared to the D39*Δply* mutant in both absence and presence of IAV infection (respectively p < 0.001 and p < 0.0001) (Figures 2A and 2B). The proportions of high-shedding events for the *Δply* mutants (exceeding 100 CFU and 2,000 CFU/tapping) were similar to those observed with the wild-type strains (Supplementary figure S2A). These results indicate that in index mice, pneumolysin did not play a significant role in nasopharyngeal shedding of pneumococci.

Transmission of pneumolysin-deficient mutants occurred only during IAV co-infection (Figure 2B), keeping similar shedding kinetics to that of the parent strains, where 100% of contact mice shed the ST217*Δply* mutant, while the D39*Δply* mutant was shed by only 67% of mice (Supplementary figure S2B). Interestingly, in contact mice, mean shedding was significantly higher with the ST217*Δply* mutant compared with the parent ST217 (p < 0.05) (Figure 2B).

No significant differences were observed in nasopharyngeal bacterial loads in index mice colonised with the *Δply* mutants compared to index mice colonised with the wild-type strains (Figure 3A). However, index mice were colonised with the D39*Δply* mutant at a significantly lower level than index mice colonised with the ST217*Δply* mutant (Figure 3A), which may have resulted in the difference of shedding between both index groups (Figure 2B). Interestingly, although contact mice that acquired pneumolysin-deficient mutants had equivalent bacterial loads in their nasopharynx compared to contact mice that acquired the parent strains (Figure 3A), both ST217- and D39-*Δply* mutants showed increased transmission rates (respectively 92% and 67%) compared with ST217 and D39 wild-type strains (respectively 58% and 24%) (Figure 3B). Altogether, these results suggest that the absence of pneumolysin had no effect on the ability of pneumococci to successfully colonise the nasopharynx of donor mice, or to be subsequently shed. Interestingly however, following transmission from donor to contact mice, the absence of pneumolysin increased the ability of the pneumococcus to disseminate within the population and colonise a significantly higher number of new hosts, perhaps indicating the lack or avoidance of a protective immune response normally triggered by the presence of pneumolysin.

### *S. pneumoniae* triggers pneumolysin-dependent macrophage recruitment following transmission

Analysis of the host immune response in the nasopharynx of mice at day 10 post infection showed that compared to the control group of mice infected with IAV alone, index and contact mice colonised with wild-type ST217 or D39 had increased numbers of macrophages (CD45^+^ CD68^+^ F4/80^+^), neutrophils (CD45^+^ Gr-1^high+^), Tbet^+^ Th1 cells, RORγt^+^ Th17 cells and FoxP3^+^ T regulatory cells, indicating that pneumococcal colonisation and acquisition stimulated the recruitment of both innate and adaptive immune cells into the nasopharynx (Supplementary figures S4A and 4B). No statistical difference was observed in numbers of host cells between ST217- and D39-colonised mice (Supplementary figures S4A and 4B), suggesting that both strains induced similar immune responses.

**Figure 4.**
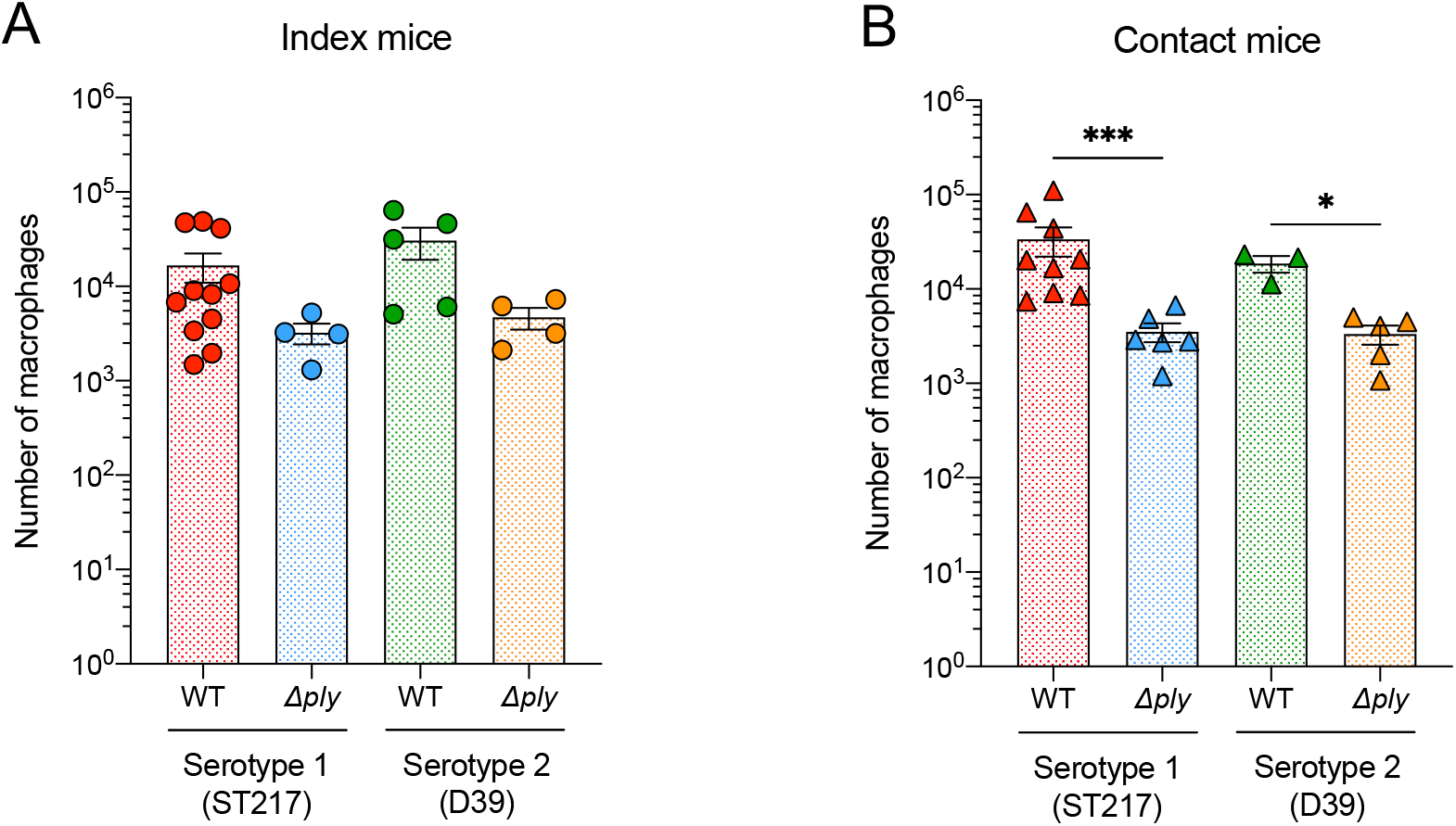
*S. pneumoniae* triggers pneumolysin-dependent macrophage recruitment in nasopharynx following transmission. Infections with *S. pneumoniae* are described in Figure 1A. At day 10 post infection, both index and contact mice colonised with serotype 1 ST217, serotype 2 (D39) or the isogenic *Δply* mutants were euthanised. Nasopharyngeal tissues were collected and processed to obtain single cell suspensions. Numbers of macrophages (CD45^+^ (FITC), CD68^+^ (PerCP/Cyanine5.5) and F4/80^+^ (Pacific Blue™) events) in nasopharynx of index (*A*) and contact (*B*) mice were determined by flow cytometry. Each symbol represents a single mouse (n = 3-11) and the mean ± SEM value is indicated. Statistical analysis was performed using the Mann-Whitney *U* test comparing mice colonised with the wild-type (WT) strains to mice colonised with the respective *Δply* mutants. *P < 0.05 and ***P < 0.001. Data represent results of multiple independent experiments (n = 1-4).

Colonisation with pneumolysin-deficient mutants also led to the recruitment of increased numbers of immune cells compared to IAV-infected control mice (Supplementary figures S3A and 3B). However, compared to mice colonised with the wild-type parent strains, index mice tended to have less macrophages in their nasopharynx when colonised with the *Δply* mutants (Figure 4A), and this decrease was significant for both pneumococcal strains in contact mice (Figure 4B), suggesting that macrophage recruitment is likely to be pneumolysin-dependent. This pneumolysin effect was not observed for any other immune cell type.

## Discussion

*S. pneumoniae* serotype 1 is the predominant cause of IPD in sub-Saharan and West Africa, despite its low carriage prevalence and short carriage duration in humans (∼ 9 days) (12, 13). Although a deeper understanding of its pathogenicity has emerged recently (8, 34, 39–41), there is a lack of knowledge of its transmission dynamics, a key factor in the emergence of disease outbreaks.

The first step for transmission is nasopharyngeal colonisation. ST217 showed a higher density of colonisation in young adult (index) mice than D39 within three days post infection, although mice were infected with an equivalent challenge dose, suggesting that ST217 had a better ability to replicate in the nasopharynx in the early stages of colonisation. Following IAV co-infection however, ST217 and D39, colonised index mice at equivalent CFU density. This is in line with our recent publication showing stable nasopharyngeal colonisation of both strains in adult Balb/c mice (34).

The second step for transmission is the ability of colonised individuals to shed pneumococci. In the absence of IAV infection in index mice, lower shedding of D39 was associated with lower colonisation compared to ST217, indicating that colonisation level may have impacted upon shedding during mono-infection. This is consistent with what has been observed in the neonate/infant mouse model, where a low level of D39-shedding was directly associated with poor colonisation of pups (42). However, as also previously observed in the infant mouse model (32, 36), co-infection with IAV increased pneumococcal shedding and transmission in our young adult mouse model as well. Indeed, increased levels of ST217-shedding in index mice occurred in the context of equivalent ST217-D39-nasopharyngeal CFU density and similar viral loads, i.e., increased ST217-shedding was not a reflection of any difference in the bacterial or viral loads between the two index groups, but a real ability of ST217 to be shed more easily, and thus, better spread and transmit to contact individuals.

Prior to the viral co-infection, no shedding events were detected in contact mice, suggesting that no transmission occurred during this period. It is likely that during infection with the pneumococcus alone, the level of shedding was not high enough in donor mice to induce transmission in contact mice. However, IAV co-infection was associated with increased CFU in index mice, resulting in increased shedding that allowed transmission of both strains to contact mice, shedding pneumococci in turn. This suggests that in our young adult model there was a shedding threshold to exceed in donor mice to induce transmission.

Serotype 1 ST217 displayed significantly higher transmission rates than serotype 2 (D39), with ST217 acquired by a significantly greater proportion of contact mice than D39 (58% versus 24% respectively). Interestingly, following its transmission, ST217 tended to colonise contact mice at a higher level compared to D39, suggesting that ST217 had a better ability to colonise a new naïve host. Recently, Abruzzo et al. showed that the likelihood for the pneumococcus to successfully establish in a new host upon its acquisition is more related to the transmission rate of each serotype rather than due to the dominance of a serotype in the upper respiratory tract of the donor host (43). This supports our data showing that despite similar nasopharyngeal densities in index mice over time, serotype 1 ST217 was much more transmissible than serotype 2 (D39) and showed a higher ability to disseminate in mice. Kono et al. have observed in infant mice that a tight bottleneck occurs in transmission following exit from a colonised host and before establishment in a naïve host, where a single bacterium would enter the new host and once established, outcompete new entering organisms (44). In our young mouse model, this would mean that a single ST217-pneumococcus may have been transmitted to contact mice and it was able to proliferate at a higher level compared to a D39-pneumococcus. Furthermore, ST217 was transmitted rapidly to naïve individuals as pneumococci were detected at very low level (1-2 CFUs) in three contact mice at days 4 and 6 post infection compared to D39 which was detected from day 7 onwards, suggesting that the window for ST217-acquisition by a new host would be short. Associated with a short carriage duration (13), our data suggest that rapid host-to-host transmission may be a driver of serotype 1 spread in the community. Although no D39-pneumococci were detected before day 7 in contact mice, we cannot exclude the possibility that mice may have acquired D39 at a non-detectable level.

Several studies have shown that pneumolysin is a key factor for pneumococcal establishment in the nasopharynx, as pneumolysin-deficient mutants display reduced colonisation (45, 46). Zafar et al., reported that pneumolysin-induced inflammation was required to promote (high) shedding in their neonate/infant mouse model as a *ply –* mutant showed significantly reduced high-shedding events in index pups under mono-infection conditions and a shedding also significantly reduced in the context of IAV co-infection (47). This contrasts with our results where *Δply* mutants in the ST217 and D39 genetic backgrounds showed equivalent or higher levels and rates of shedding compared to the parent strains, indicating that in young adult mice, pneumolysin deficiency did not reduce shedding during pneumococcal mono-infection or co-infection with IAV. In addition, pneumolysin-deficient mutants colonised mice at similar levels to those of the wild-type isolates, indicating that in our model at least, pneumolysin *did not play* an essential role in colonisation. Our data show how the models used, *young adult* versus *neonate*/*infant*, may impact directly on the role of pneumolysin in colonisation and transmission. Human and murine infants are known to have a limited B cell response due to low expression of co-receptors and strongly lack Th1 responses limiting their capacity to respond to foreign antigens (5, 48, 49). This may explain why infant mice are more susceptible to the effects of pneumolysin during transmission, as the toxin promotes mucosal inflammation in the nasopharynx which aids transmission between pups (47). In contrast, older children and young adults who are immunologically competent, may better control pneumolysin driven inflammation through CD4^+^ T cell and Treg cell control (37), and our transmission model in young adult mouse would better reflects these conditions.

Zafar et al. also suggested that pneumolysin was no longer required for transmission during viral co-infection, as in absence of the toxin, their *ply –* mutant was still transmitted in IAV co-infected pups, the virus promoting nasopharyngeal inflammation necessary for transmission (47). Interestingly, we observed increased transmission rates for the *Δply* mutants, although contact mice were colonised with WT strains or pneumolysin-deficient mutants at similar levels (Figure 3), suggesting that without the well established pro-inflammatory and immune cell activating effects of pneumolysin, the pneumococcus had a better capacity to disseminate within the population, and be transmitted to and colonise a greater number of individuals. This is supported by our data which showed significantly reduced macrophage numbers in the nasopharynx of contact mice following pneumolysin-deficient pneumococcal acquisition, indicating that recruitment of macrophages is pneumolysin dependent and that the reduction of macrophage influx observed may have been critical enough to increase susceptibility of recipient animals to pneumococcal acquisition and impair bacterial clearance which promoted establishment of the pneumococcus in new hosts. This pneumolysin-dependent macrophage recruitment was also evident in index mice albeit not statistically significant. Our data are consistent with previous studies reporting the important role of pneumolysin in macrophage activation and recruitment (50, 51) and the key role of macrophages in phagocytic protection against pneumococcal carriage and infection (37, 52, 53). Pneumolysin has also been demonstrated to interact directly with various cells of the host innate immune system and indirectly by driving inflammation, such as activation of the NLRP3 inflammasome in macrophages and dendritic cells, mediating production of pro-inflammatory cytokines such as IL-1β in response to infection (54–56).

We also showed that colonisation with *S. pneumoniae* resulted in significantly increased recruitment of adaptive immune cells (Th17, Th1 and T regulatory cells) into the nasopharynx, which is consistent with previous studies highlighting the role of CD4^+^ T cells in the host immune response to pneumococcal infection in mice and humans (57–62). Pneumolysin-deficient mutants induced similar host cell recruitment to their parent strains, suggesting that the absence of pneumolysin did not significantly impact the immune response (except for macrophages) during pneumococcal colonisation in our young adult model.

Our study offers new insights into the understanding of transmission of a hypervirulent pneumococcal strain during host-to-host transmission. Our results demonstrate that high shedding and transmission rate rather than carriage rate (or duration) may explain why serotype 1 ST217 exhibits high prevalence in IPD outbreaks and that strategies to prevent host-to-host transmission must be considered to reduce serotype 1 associated pneumococcal disease within populations.

## Supporting information

Baltazar et al_Supplementary data

## Acknowledgements

This work was supported by the Joint Programming Initiative on Antimicrobial Resistance (JPIAMR) and the UK Medical Research Council Programme Grant (MR/P011284/1) awarded to A.K. and a Mahidol – Liverpool University funded PhD studentship to T.A.

We are grateful to Professor James Stewart for providing the influenza virus A/HKx31 (H3N2) and the Biomedical Services Unit of the University of Liverpool.

## Supplementary figures

**Supplementary figure S1. Influenza A virus increases pneumococcal colonisation in nasopharynx of young adult mice.**

Colonisation densities of serotype 1 ST217 and serotype 2 (D39) in the nasopharynx of index mice at day 10 post infection during IAV co-infection conditions. Each symbol represents a single mouse and the mean ± SEM value is indicated. Statistical analysis was performed using the Mann-Whitney *U* test. ****P < 0.0001. Data represent results of multiple independent experiments (n = 1-5).

**Supplementary figure S2. Serotype 1 ST217 displays high rates of shedding.**

Rates of shedding of serotype 1 ST217, serotype 2 (D39) and the isogenic *Δply* mutants in index *(A)* and contact *(B)* mice. Statistical analysis was performed using the Fisher’s exact test. ^a^ in comparison to ST217; ^b^ in comparison to wild-type strains. Data represent results of multiple independent experiments (n = 1-5).

**Supplementary figure S3. Serotype 1 ST217 produces a thicker capsule than serotype 2 (D39).**

Comparison of the capsule size between serotype 1 ST217 and serotype 2 (D39). The capsule size was determined as the width of bacteria excluded by FITC-dextran. Each symbol represents the calculated mean area per bacterium (in square pixels). Each strain was analysed in duplicate or triplicate in 10-11 independent experiments performed on different days. The mean ± SD value is indicated. Statistical analysis was performed using the Mann-Whitney *U* test. ****P < 0.0001.

**Supplementary figure S4. Host immune response in the nasopharynx during pneumococcal colonisation and following transmission.**

Infections with *S. pneumoniae* are described in Figure 1A. In the control group “IAV alone”, young adult mice were intranasally inoculated with PBS on day zero. Three days after mock inoculation, all mice were intranasally infected with influenza virus A/HKx31 (X31, H3N2). At day 10 post infection, both index and contact mice colonised with serotype 1 ST217, serotype 2 (D39) or the isogenic *Δply* mutants, and control mice were euthanised. Nasopharyngeal tissues were collected and processed to obtain single cell suspensions. Numbers of macrophages (CD45^+^ (FITC), CD68^+^ (PerCP/Cyanine5.5) and F4/80^+^ (Pacific Blue™) events), numbers of neutrophils (CD45^+^ (FITC) and Gr-1^high+^ (APC/Cyanine7) events), numbers of Th1 CD4^+^ T cells (CD45^+^ (FITC), CD4^+^ (APC/Cy7) and T-bet^+^ (APC) events), numbers of Th17 CD4^+^ T cells (CD45^+^ (FITC), CD4^+^ (APC/Cy7) and RORγt^+^ (PerCP-Cy™5.5) events) and numbers of T regulatory (Treg) cells (CD45^+^ (FITC), CD4^+^ (APC/Cy7) and FoxP3^+^ (PE) events) in nasopharynx of index (*A*) and contact (*B*) mice were determined by flow cytometry. Each symbol represents a single mouse (n = 3-11) and the mean ± SEM value is indicated. Statistical analysis was performed using the Kruskal-Wallis test with Dunn’s post test. *P < 0.05, **P < 0.01 and ***P < 0.001. Data represent results of multiple independent experiments (n = 1-4).

## Supplementary tables

**Supplementary table S1. Bacterial strains used in this study.**

**Supplementary table S2. Monoclonal antibodies panels and dilution used for flow cytometry analysis.**

## References

1. Webster LT, Hughes TP. The epidemiology of pneumococcus infection: the incidence and spread of pneumococci in the nasal passages and throats of healthy persons. J Exp Med 1931;53:535–552.

2. Bogaert D, De Groot R, Hermans PWM. *Streptococcus pneumoniae* colonisation: the key to pneumococcal disease. Lancet Infect Dis 2004;4:144–154.

3. Marks LR, Davidson BA, Knight PR, Hakansson AP. Interkingdom signaling induces *Streptococcus pneumoniae* biofilm dispersion and transition from asymptomatic colonization to disease. mBio 2013;4:e00438–13.

4. Brown JM, Hammerschmidt S, Orihuela C. Streptococcus Pneumoniae: molecular mechanisms of host-pathogen interactions. Academic Press; 2015.

5. Brooks LRK, Mias GI. *Streptococcus pneumoniae*’s virulence and host immunity: aging, diagnostics, and prevention. Front Immunol 2018;9:1366.

6. Antonio M, Hakeem I, Awine T, Secka O, Sankareh K, Nsekpong D, et al. Seasonality and outbreak of a predominant *Streptococcus pneumoniae* serotype 1 clone from The Gambia: expansion of ST217 hypervirulent clonal complex in West Africa. BMC Microbiol 2008;8:198.

7. Blumental S, Moïsi JC, Roalfe L, Zancolli M, Johnson M, Burbidge P, et al. *Streptococcus pneumoniae* Serotype 1 burden in the African meningitis belt: exploration of functionality in specific antibodies. Clin Vaccine Immunol 2015;22:404–412.

8. Chaguza C, Yang M, Jacques LC, Bentley SD, Kadioglu A. Serotype 1 pneumococcus: epidemiology, genomics, and disease mechanisms. Trends Microbiol 2022;30:581–592.

9. Gessner BD, Mueller JE, Yaro S. African meningitis belt pneumococcal disease epidemiology indicates a need for an effective serotype 1 containing vaccine, including for older children and adults. BMC Infect Dis 2010;10:22.

10. Kwambana-Adams BA, Asiedu-Bekoe F, Sarkodie B, Afreh OK, Kuma GK, Owusu-Okyere G, et al. An outbreak of pneumococcal meningitis among older children (≥5 years) and adults after the implementation of an infant vaccination programme with the 13-valent pneumococcal conjugate vaccine in Ghana. BMC Infect Dis 2016;16:575.

11. Leimkugel J, Adams Forgor A, Gagneux S, Pflüger V, Flierl C, Awine E, et al. An Outbreak of serotype 1 *Streptococcus pneumoniae* meningitis in Northern Ghana with features that are characteristic of *Neisseria meningitidis* meningitis epidemics. J Infect Dis 2005;192:192–199.

12. Ebruke C, Roca A, Egere U, Darboe O, Hill PC, Greenwood B, et al. Temporal changes in nasopharyngeal carriage of *Streptococcus pneumoniae* serotype 1 genotypes in healthy Gambians before and after the 7-valent pneumococcal conjugate vaccine. PeerJ 2015;3:e903.

13. Abdullahi O, Karani A, Tigoi CC, Mugo D, Kungu S, Wanjiru E, et al. Rates of acquisition and clearance of pneumococcal serotypes in the nasopharynges of children in Kilifi District, Kenya. J Infect Dis 2012;206:1020–1029.

14. Ritchie ND, Mitchell TJ, Evans TJ. What is different about serotype 1 pneumococci? Future Microbiol 2012;7:33–46.

15. von Mollendorf C, Cohen C, Tempia S, Meiring S, de Gouveia L, Quan V, et al. Epidemiology of serotype 1 invasive pneumococcal disease, South Africa, 2003–2013. Emerg Infect Dis 2016;22:261–270.

16. Yaro S, Lourd M, Traoré Y, Njanpop-Lafourcade B-M, Sawadogo A, Sangare L, et al. Epidemiological and molecular characteristics of a highly lethal pneumococcal meningitis epidemic in Burkina Faso. Clin Infect Dis 2006;43:693–700.

17. du Plessis M, Allam M, Tempia S, Wolter N, de Gouveia L, von Mollendorf C, et al. Phylogenetic analysis of invasive serotype 1 pneumococcus in South Africa, 1989 to 2013. J Clin Microbiol 2016;54:1326–1334.

18. Donkor ES. Understanding the pneumococcus: transmission and evolution. Front Cell Infect Microbiol 2013;3:7.

19. Musher DM. How contagious are common respiratory tract infections? N Engl J Med 2003;348:1256–1266.

20. Walsh RL, Camilli A. *Streptococcus pneumoniae* is desiccation tolerant and infectious upon rehydration. mBio 2011;2:e00092–11.

21. Marks LR, Reddinger RM, Hakansson AP. Biofilm formation enhances fomite survival of *Streptococcus pneumoniae* and *Streptococcus pyogenes*. Infect Immun 2014;82:1141–1146.

22. Numminen E, Chewapreecha C, Turner C, Goldblatt D, Nosten F, Bentley SD, et al. Climate induces seasonality in pneumococcal transmission. Sci Rep 2015;5:11344.

23. Domenech de Cellès M, Arduin H, Lévy-Bruhl D, Georges S, Souty C, Guillemot D, et al. Unraveling the seasonal epidemiology of pneumococcus. Proc Natl Acad Sci U S A 2019;116:1802–1807.

24. Hodges RG, MacLeod CM. Epidemic pneumococcal pneumonia; the influence of population characteristics and environment. Am J Hyg 1946;44:193–206.

25. Bosch AATM, Biesbroek G, Trzcinski K, Sanders EAM, Bogaert D. Viral and bacterial interactions in the upper respiratory tract. PLOS Pathog 2013;9:e1003057.

26. Grijalva CG, Griffin MR, Edwards KM, Williams JV, Gil AI, Verastegui H, et al. The role of influenza and parainfluenza infections in nasopharyngeal pneumococcal acquisition among young children. Clin Infect Dis 2014;58:1369–1376.

27. Karppinen S, Teräsjärvi J, Auranen K, Schuez-Havupalo L, Siira L, He Q, et al. Acquisition and transmission of *Streptococcus pneumoniae* are facilitated during rhinovirus infection in families with children. Am J Respir Crit Care Med 2017;196:1172–1180.

28. McCullers JA. Insights into the interaction between influenza virus and pneumococcus. Clin Microbiol Rev 2006;19:571–582.

29. Horácio AN, Silva-Costa C, Lopes JP, Ramirez M, Melo-Cristino J, Portuguese Group for the Study of Streptococcal Infections. Serotype 3 remains the leading cause of invasive pneumococcal disease in adults in Portugal (2012-2014) despite continued reductions in other 13-valent conjugate vaccine serotypes. Front Microbiol 2016;7:1616.

30. Imöhl M, Reinert RR, Ocklenburg C, Linden M van der. Association of serotypes of *Streptococcus pneumoniae* with age in invasive pneumococcal disease. J Clin Microbiol 2010;48:1291–1296.

31. McCullers JA, McAuley JL, Browall S, Iverson AR, Boyd KL, Normark BH. Influenza enhances susceptibility to natural acquisition and disease from *Streptococcus pneumoniae* in ferrets. J Infect Dis 2010;202:1287–1295.

32. Diavatopoulos DA, Short KR, Price JT, Wilksch JJ, Brown LE, Briles DE, et al. Influenza A virus facilitates *Streptococcus pneumoniae* transmission and disease. FASEB J 2010;24:1789–1798.

33. Malley R, Stack AM, Ferretti ML, Thompson CM, Saladino RA. Anticapsular polysaccharide antibodies and nasopharyngeal colonization with *Streptococcus pneumoniae* in infant rats. J Infect Dis 1998;178:878–882.

34. Jacques LC, Panagiotou S, Baltazar M, Senghore M, Khandaker S, Xu R, et al. Increased pathogenicity of pneumococcal serotype 1 is driven by rapid autolysis and release of pneumolysin. Nat Commun 2020;11:1–13.

35. Gates MA, Thorkildson P, Kozel TR. Molecular architecture of the *Cryptococcus neoformans* capsule. Mol Microbiol 2004;52:13–24.

36. Richard AL, Siegel SJ, Erikson J, Weiser JN. TLR2 signaling decreases transmission of *Streptococcus pneumoniae* by limiting bacterial shedding in an infant mouse influenza A co-infection model. PLOS Pathog 2014;10:e1004339.

37. Neill DR, Coward WR, Gritzfeld JF, Richards L, Garcia-Garcia FJ, Dotor J, et al. Density and duration of pneumococcal carriage is maintained by Transforming Growth Factor β1 and T regulatory cells. Am J Respir Crit Care Med 2014;189:1250–1259.

38. Zafar MA, Hamaguchi S, Zangari T, Cammer M, Weiser JN. Capsule type and amount affect shedding and transmission of *Streptococcus pneumoniae*. mBio 2017;8:e00989–17.

39. Bricio-Moreno L, Ebruke C, Chaguza C, Cornick J, Kwambana-Adams B, Yang M, et al. Comparative genomic analysis and in vivo modeling of *Streptococcus pneumoniae* ST3081 and ST618 isolates reveal key genetic and phenotypic differences contributing to clonal replacement of serotype 1 in The Gambia. J Infect Dis 2017;216:1318–1327.

40. Cornick JE, Tastan Bishop Ö, Yalcin F, Kiran AM, Kumwenda B, Chaguza C, et al. The global distribution and diversity of protein vaccine candidate antigens in the highly virulent *Streptococcus pneumoniae* serotype 1. Vaccine 2017;35:972–980.

41. Bricio-Moreno L, Chaguza C, Yahya R, Shears RK, Cornick JE, Hokamp K, et al. Lower density and shorter duration of nasopharyngeal carriage by pneumococcal serotype 1 (ST217) may explain its increased invasiveness over other serotypes. mBio 2020;11:e00814–20.

42. Zafar MA, Kono M, Wang Y, Zangari T, Weiser JN. Infant mouse model for the study of shedding and transmission during *Streptococcus pneumoniae* monoinfection. Infect Immun 2016;84:2714–2722.

43. Abruzzo AR, Aggarwal SD, Sharp ME, Bee GCW, Weiser JN. Serotype-dependent effects on the dynamics of pneumococcal colonization and implications for transmission. mBio 2022;13:e00158–22.

44. Kono M, Zafar MA, Zuniga M, Roche AM, Hamaguchi S, Weiser JN. Single cell bottlenecks in the pathogenesis of *Streptococcus pneumoniae*. PLOS Pathog 2016;12:e1005887.

45. Kadioglu A, Taylor S, Iannelli F, Pozzi G, Mitchell TJ, Andrew PW. Upper and lower respiratory tract infection by *Streptococcus pneumoniae* is affected by pneumolysin deficiency and differences in capsule type. Infect Immun 2002;70:2886–2890.

46. Richards L, Ferreira DM, Miyaji EN, Andrew PW, Kadioglu A. The immunising effect of pneumococcal nasopharyngeal colonisation; protection against future colonisation and fatal invasive disease. Immunobiology 2010;215:251–263.

47. Zafar MA, Wang Y, Hamaguchi S, Weiser JN. Host-to-host transmission of *Streptococcus pneumoniae* is driven by its inflammatory toxin, pneumolysin. Cell Host Microbe 2017;21:73–83.

48. Basha S, Surendran N, Pichichero M. Immune responses in neonates. Expert Rev Clin Immunol 2014;10:1171–1184.

49. Simon AK, Hollander GA, McMichael A. Evolution of the immune system in humans from infancy to old age. Proc R Soc B Biol Sci 2015;282:20143085.

50. Bewley MA, Naughton M, Preston J, Mitchell A, Holmes A, Marriott HM, et al. Pneumolysin activates macrophage lysosomal membrane permeabilization and executes apoptosis by distinct mechanisms without membrane pore formation. mBio 2014;5:e01710–14.

51. Das R, LaRose MI, Hergott CB, Leng L, Bucala R, Weiser JN. Macrophage migration inhibitory factor promotes clearance of pneumococcal colonization. J Immunol 2014;193:764–772.

52. Jochems SP, Weiser JN, Malley R, Ferreira DM. The immunological mechanisms that control pneumococcal carriage. PLOS Pathog 2017;13:e1006665.

53. Knapp S, Leemans JC, Florquin S, Branger J, Maris NA, Pater J, et al. Alveolar macrophages have a protective antiinflammatory role during murine pneumococcal pneumonia. Am J Respir Crit Care Med 2003;167:171–179.

54. Harvey RM, Hughes CE, Paton AW, Trappetti C, Tweten RK, Paton JC. The impact of pneumolysin on the macrophage response to *Streptococcus pneumoniae* is strain-dependent. PLOS ONE 2014;9:e103625.

55. Lemon JK, Miller MR, Weiser JN. Sensing of interleukin-1 cytokines during *Streptococcus pneumoniae* colonization contributes to macrophage recruitment and bacterial clearance. Infect Immun 2015;83:3204–3212.

56. Surabhi S, Cuypers F, Hammerschmidt S, Siemens N. The role of NLRP3 inflammasome in pneumococcal infections. Front Immunol 2020;11:614801.

57. Kadioglu A, Coward W, Colston MJ, Hewitt CRA, Andrew PW. CD4-T-lymphocyte interactions with pneumolysin and pneumococci suggest a crucial protective role in the host response to pneumococcal infection. Infect Immun 2004;72:2689–2697.

58. Kemp K, Bruunsgaard H, Skinhøj P, Pedersen BK. Pneumococcal infections in humans are associated with increased apoptosis and trafficking of type 1 cytokine-producing T cells. Infect Immun 2002;70:5019–5025.

59. Malley R, Trzcinski K, Srivastava A, Thompson CM, Anderson PW, Lipsitch M. CD4+ T cells mediate antibody-independent acquired immunity to pneumococcal colonization. Proc Natl Acad Sci 2005;102:4848–4853.

60. Olliver M, Hiew J, Mellroth P, Henriques-Normark B, Bergman P. Human monocytes promote Th1 and Th17 responses to *Streptococcus pneumoniae*. Infect Immun 2011;79:4210–4217.

61. van Rossum AMC, Lysenko ES, Weiser JN. Host and bacterial factors contributing to the clearance of colonization by *Streptococcus pneumoniae* in a murine model. Infect Immun 2005;73:7718–7726.

62. Zhang Q, Bagrade L, Bernatoniene J, Clarke E, Paton JC, Mitchell TJ, et al. Low CD4 T cell immunity to pneumolysin is associated with nasopharyngeal carriage of pneumococci in children. J Infect Dis 2007;195:1194–1202.

63. Lacks S, Hotchkiss RD. Formation of amylomaltase after genetic transformation of pneumococcus. Biochim Biophys Acta 1960;45:155–163.

