## Supplementary material for "Hypervirulent pneumococci display high levels of nasopharyngeal shedding and rapid onward transmission": Baltazar et al_Supplementary data

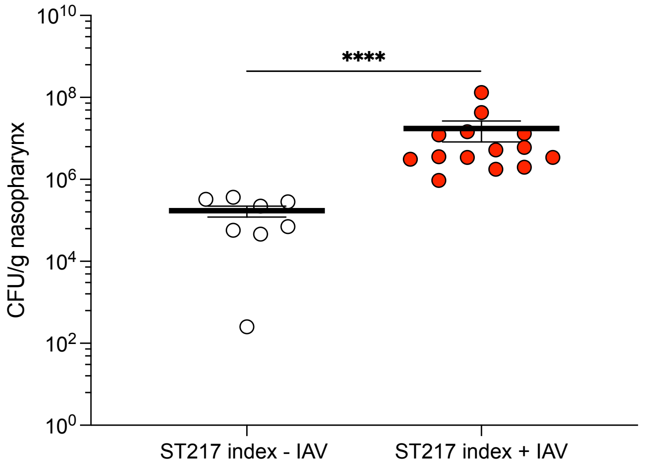


**Supplementary figure S1. Influenza A virus increases pneumococcal colonisation in nasopharynx of young adult mice.**
Colonisation densities of serotype 1 ST217 and serotype 2 (D39) in the nasopharynx of index mice at day 10 post infection during IAV co-infection conditions. Each symbol represents a single mouse and the mean ± SEM value is indicated. Statistical analysis was performed using the Mann-Whitney *U* test. ****P < 0.0001. Data represent results of multiple independent experiments (n = 1-5).

A

| Conditions | Rates of shedding in index mice | ST217 | D39 | ST217 *Δply* | D39 *Δply* |
|---|---|---|---|---|---|
| - IAV | > 100 CFU/tapping/ Total shedding events | 29/54 | 1/44 | 10/24 | 1/23 |
| | Shedding > 100 CFU (%) | 54 | 2 | 42 | 4 |
| | P-value | — | < 0.0001[a] | 0.4622[b] | > 0.9999[b] |
| + IAV | > 2,000 CFU/tapping/ Total shedding events | 28/92 | 7/86 | 22/48 | 5/42 |
| | Shedding > 2,000 CFU (%) | 30 | 8 | 46 | 12 |
| | P-value | — | 0.0003[a] | 0.0943[b] | 0.5272[b] |

B

| Condition | Rates of shedding in contact mice | ST217 | D39 | ST217 *Δply* | D39 *Δply* |
|---|---|---|---|---|---|
| + IAV | Number of contact mice shedding/Total number of contact mice | 22/24 | 9/21 | 12/12 | 8/12 |
| | Shedding (%) | 92 | 43 | 100 | 67 |
| | P-value | — | 0.0008[a] | 0.5429[b] | 0.2818[b] |

**Supplementary figure S2. Serotype 1 ST217 displays high rates of shedding.**
Rates of shedding of serotype 1 ST217, serotype 2 (D39) and the isogenic *Δply* mutants in index *(A)* and contact *(B)* mice. Statistical analysis was performed using the Fisher's exact test. [a] in comparison to ST217; [b] in comparison to wild-type strains. Data represent results of multiple independent experiments (n = 1-5).

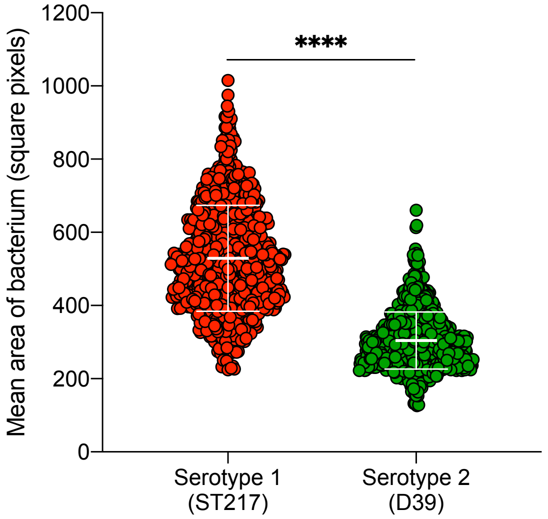


**Supplementary figure S3. Serotype 1 ST217 produces a thicker capsule than serotype 2 (D39).**

Comparison of the capsule size between serotype 1 ST217 and serotype 2 (D39). The capsule size was determined as the width of bacteria excluded by FITC-dextran. Each symbol represents the calculated mean area per bacterium (in square pixels). Each strain was analysed in duplicate or triplicate in 10-11 independent experiments performed on different days. The mean ± SD value is indicated. Statistical analysis was performed using the Mann-Whitney *U* test. ****$P < 0.0001$.

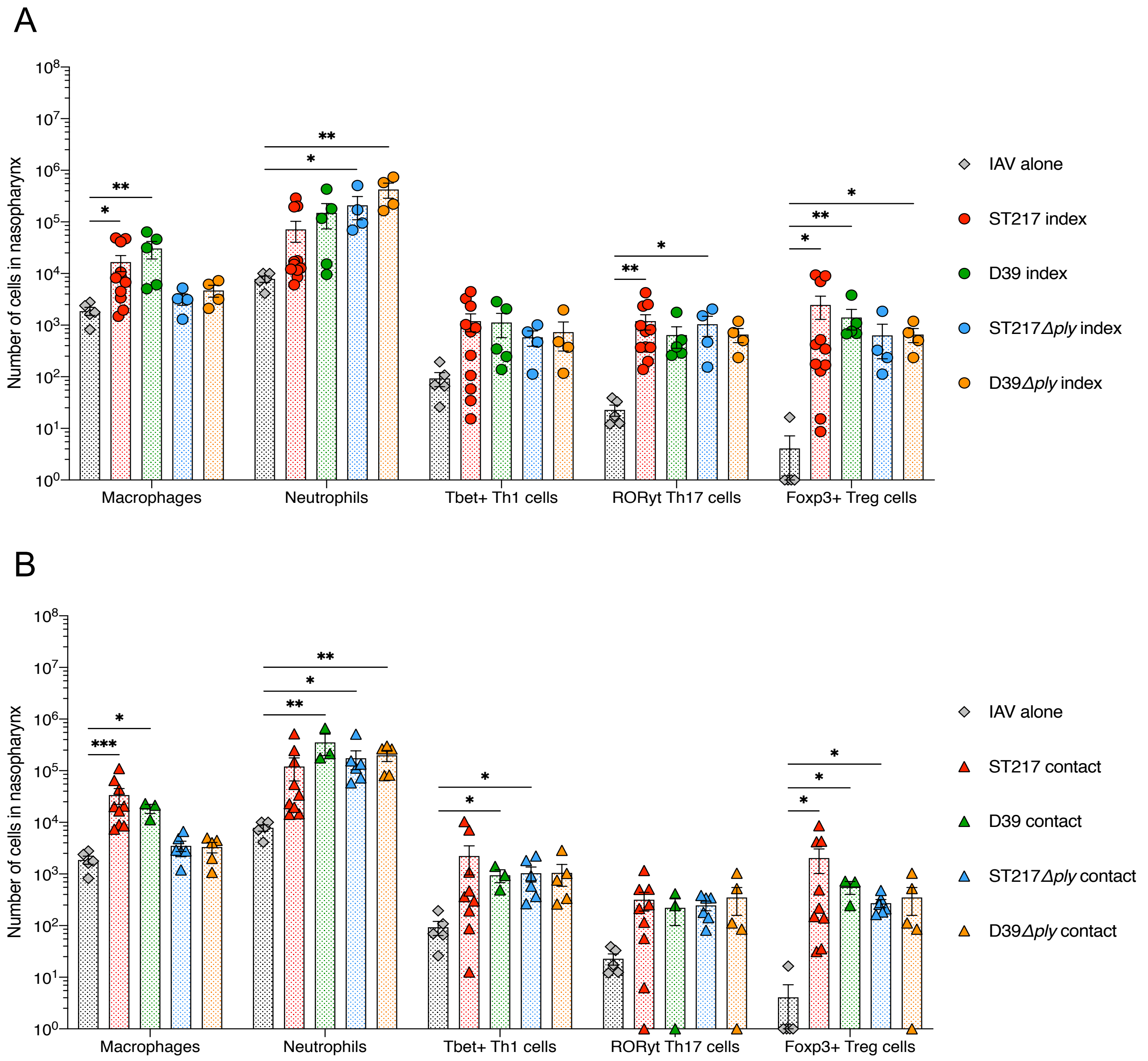


**Supplementary figure S4. Host immune response in the nasopharynx during pneumococcal colonisation and following transmission.**
Infections with *S. pneumoniae* are described in Figure 1A. In the control group "IAV alone", young adult mice were intranasally inoculated with PBS on day zero. Three days after mock inoculation, all mice were intranasally infected with influenza virus A/HKx31 (X31, H3N2). At day 10 post infection, both index and contact mice colonised with serotype 1 ST217, serotype 2 (D39) or the isogenic *Δply* mutants, and control mice were euthanised. Nasopharyngeal tissues were collected and processed to obtain single cell suspensions. Numbers of macrophages ($CD45^+$ (FITC), $CD68^+$ (PerCP/Cyanine5.5) and $F4/80^+$ (Pacific Blue™) events), numbers of neutrophils ($CD45^+$ (FITC) and $Gr\text{-}1^{high+}$ (APC/Cyanine7) events), numbers of Th1 $CD4^+$ T cells ($CD45^+$ (FITC), $CD4^+$ (APC/Cy7) and $T\text{-}bet^+$ (APC) events), numbers of Th17 $CD4^+$ T cells ($CD45^+$ (FITC), $CD4^+$ (APC/Cy7) and $ROR\gamma t^+$ (PerCP-Cy™5.5) events) and numbers of T regulatory (Treg) cells ($CD45^+$ (FITC), $CD4^+$ (APC/Cy7) and $FoxP3^+$ (PE) events) in nasopharynx of index (*A*) and contact (*B*) mice were determined by flow cytometry. Each symbol represents a single mouse (n = 3-11) and the mean ± SEM value is indicated. Statistical analysis was performed using the Kruskal-Wallis test with Dunn's post test. *P < 0.05, **P < 0.01 and ***P < 0.001. Data represent results of multiple independent experiments (n = 1-4).

**Supplementary Table S1. Strains used in this study.**

| ***Streptococcus pneumoniae* strains** | **Description** | **References** |
|---|---|---|
| Serotype 1 ST217 | Invasive clinical isolate isolated from the cerebrospinal fluid of a male adult patient diagnosed with pneumococcal meningitis and hospitalised at the Queen Elisabeth Hospital, in Blantyre, Malawi. | This study. |
| Serotype 1 ST217*Δply* | Isogenic pneumolysin-deficient mutant of serotype 1 ST217 in which *ply* gene was replaced by the *aphA3* gene conferring resistance to kanamycin. | This study. |
| Serotype 2 strain D39 (NCTC 7466) | Obtained from the National Collection of Type Cultures, London, United Kingdom. | |
| Serotype 2 strain D39*Δply* | Isogenic pneumolysin-deficient mutant of serotype 2 strain D39 carrying an insertion-duplication mutation which interrupts the pneumolysin-coding sequence. | (1) |

Reference

1. Berry AM, Yother J, Briles DE, Hansman D, Paton JC. Reduced virulence of a defined pneumolysin-negative mutant of *Streptococcus pneumoniae*. *Infect Immun* 1989;57:2037–2042.

**Supplementary Table S2. Monoclonal antibodies and dilutions used for flow cytometry analysis.**

| Target Cell | Supplier | Antibodies used (Catalogue No.) | Host Specie | Clone | Dilution |
|---|---|---|---|---|---|
| **Neutrophils** | BioLegend® | CD45-FITC (103108) | Rat | 30-F11 | 1/400 |
| | BioLegend® | Ly-6G/Ly-6C (GR-1)-APC/Cyanine7 (108423) | Rat | RB6-8C5 | 1/400 |
| **Macrophages** | BioLegend® | CD45-FITC** | Rat | 30-F11 | 1/400 |
| | BioLegend® | CD68-PerCP/Cyanine5.5 (137009) | Rat | FA-11 | 1/400 |
| | BioLegend® | F4/80-Pacific Blue™ (123123) | Rat | BM8 | 1/300 |
| **Th1 cells** | BioLegend® | CD45-FITC** | Rat | 30-F11 | 1/400 |
| | BioLegend® | CD4-APC/Cy7 (100525) | Rat | RM4-5 | 1/400 |
| | BioLegend® | T-bet-APC (644813) | Mouse | 4B10 | 1/400 |
| **Th17 cells** | BioLegend® | CD45-FITC** | Rat | 30-F11 | 1/400 |
| | BioLegend® | CD4-APC/Cy7** | Rat | RM4-5 | 1/400 |
| | BD Biosciences | RORγt-PerCP-Cy™5.5 (562683) | Mouse | Q31-378 | 1/400 |
| **T regulatory cells** | BioLegend® | CD45-FITC** | Rat | 30-F11 | 1/400 |
| | BioLegend® | CD4-APC/Cy7** | Rat | RM4-5 | 1/400 |
| | BioLegend® | FOXP3-PE (126403) | Rat | MF-14 | 1/400 |

**: same antibody as previously described in the table.
